# Structural Basis of Enhanced Facilitated Diffusion of DNA Binding Proteins in Crowded Cellular Milieu

**DOI:** 10.1101/701557

**Authors:** P. Dey, A. Bhattacherjee

**Affiliations:** Jawaharlal Nehru University

## Abstract

DNA binding proteins (DBPs) rapidly recognize and specifically associate with their target DNA sites inside cell nucleus that contains up to 400 g/L macromolecules, most of which are proteins. While the fast association between DBPs and DNA is explained by a facilitated diffusion mechanism, where DBPs adopt a weighted combination of 3D diffusion and 1D sliding and hopping modes of transportation, the role of cellular environment that contains many nonspecifically interacting proteins and other biomolecules is mostly overlooked. By performing large scale computational simulations with an appropriately tuned model of protein and DNA in the presence of nonspecifically interacting bulk and DNA bound crowders (genomic crowders), we demonstrate the structural basis of the enhanced facilitated diffusion of DBPs inside a crowded cellular milieu through novel 1D scanning mechanisms. In the presence of bulk crowders, we identify the protein to float along the DNA under the influence of protein-crowder nonspecific interactions. The search mode is distinctly different compared to usual 1D sliding and hopping dynamics where protein diffusion is regulated by the DNA electrostatics. In contrast, the presence of genomic crowders expedite the target search process by transporting the protein over DNA segments through the formation of a transient protein-crowder bridged complex. By analyzing the ruggedness of the associated potential energy landscape, we underpin the molecular origin of the kinetic advantages of these search modes and show that they successfully explain the experimentally observed acceleration of facilitated diffusion of DBPs by molecular crowding agents and crowder concentration dependent enzymatic activity of transcription factors. Our findings provide crucial insights into gene regulation kinetics inside the crowded cellular milieu.

**SIGNIFICANCE:** 10-40% of the intracellular volume is occupied by proteins, and other biomolecules, collectively known as macromolecular crowders. Their presence has been found to promote faster translocation of DNA binding proteins (DBPs) during the search of their target DNA sites for crucial cellular processes. Using molecular simulations, we probe the underlying structural basis and underscore the existence of novel DNA scanning mechanisms actuated by protein-crowder nonspecific interactions. We show that the observed search modes are kinetically beneficial and can successfully explain the acceleration of facilitated diffusion of DBPs by molecular crowding agents and crowderconcentration dependent enzymatic activity of transcription factors.Our study sheds new light on the long-standing facilitated diffusion problem of DBPs in the crowded cellular environment for regulating gene expression.

## INTRODUCTION

The rapid and efficient association of DNA Binding Proteins (DBPs) to their target sites on genomic DNA fuels crucial biological processes such as gene expression, transcription, DNA damage repair etc. (1–3). The fast kinetics is achieved through a weighted proportion of one and three-dimensional diffusion (4, 5) of the DBPs. During 1D diffusion, proteins either slide along the helical pitch of the DNA or perform series of dissociation and re-association events of short life spans (‘hopping’) along the DNA contour. Furthermore, the proteins facilitate the search process via inter-segmental ‘jumps’(6, 7) between nearby DNA segments and are thereby able to bypass the scanning of several DNA bases. Flanking DNA sequence around the target DNA site has also been found to modulate the search process by altering the funnel-shaped protein-DNA binding energy landscape(8). These multifaceted search mechanisms have been captured in vitro via different spectroscopic approaches(9–13), and also through in silico models and simulations(14–23) at single molecule level. The in vivo condition is, however, entirely different due to the presence of an astounding concentration (~100-300 mg/ml) of dissolved macromolecules(24, 25). This means 10-40% of the total intracellular volume is occupied by the macromolecules that cause ~5-10 times high viscosity (*η*) in the cellular medium compared to the laboratory buffer solutions. Furthermore, most of these macromolecules are proteins that exert a complex variety of effects on other proteins including DBPs, through nonspecific interactions such as hydrogen bonding, ionic interactions etc. Both the factors are seemingly unfavorable for free diffusion of DBPs inside a crowded nuclear environment.

Recent experiments(26–28), however, suggest faster diffusion and enhanced enzyme activity of DNA binding proteins in the presence of molecular crowders. For example, in vitro studies on the hydrolysis of DNA by endonucleases DNase I and S1 nuclease increase substantially in a medium crowded by polyethylene glycol (PEG)(29). Facilitated diffusion in crowded medium has been observed experimentally for other systems as well, including translocation of human DNA glycosylases inside highly dense PEG solution(27), and optimized regulation kinetics of lac repressor in E. coli(30). Despite its widespread importance across different types of regulatory processes, how the crowded cellular medium enhances the facilitated diffusion of DBPs remains unclear(31–33).

Early studies proposed repeated collisions between the molecules in presence of crowding agents as the molecular basis of tighter binding of protein-DNA complexes(34, 35) that can significantly impacts the gene expression(36). Further systematic investigation of diffusion in crowded media(22, 37) supported the viewpoint and predicted that the role of bulk crowders is to prevent the dissociation of DBPs from the DNA surface and thereby promoting 1D diffusion of the searching protein along the DNA contour over 3D diffusion(38, 39). In contrast, previous studies indicate that purely repulsive crowders play a role to decouple the searching protein from the crowding agents during the 1D search regime on a linear DNA stretch(40, 41). The observation is along the line of Asakura-Oosawa model (AOM)(42) of crowder action, where the crowding agents, due to excluded volume interactions with the DNA molecule result in a preferentially depleted volume around the DNA interface. Since the region is devoid of crowder molecules, the nonspecific rotational and translational diffusion of the searching protein inside this region were found to be independent of the concentration of bulk crowders(40). The protein effectively experiences a constant microviscosity much lower than that of the bulk, which could promote faster diffusion of the protein. The model, however, is inconsistent with the protein-based cytoplasmic crowders(43–46) that exert nonspecific attractive forces on other macromolecules beside excluding the available volume for them. To this end, one should also note the experiments that have reported enhanced facilitated diffusion in the presence of synthetic crowding agents such as PEG and explained the observations exclusively on the basis of ‘volume exclusions’ by the crowding agents, misinterpreted the ‘inert’ character of the crowding agents. Indeed, recent calorimetric experiment indicates that the commonly used synthetic agents such as Dextran, glucose and PEG can significantly contribute to the enthalpic stabilization and the entropic destabilization of proteins(47). The observations and the presence of biomolecules in in vivo conditions that can mediate attractive nonspecific interactions with the DBPs necessitate probing the role of an interacting crowded environment in the facilitated diffusion of DBPs.

In this paper, we, therefore, probe the target search dynamics of DBPs in the presence of explicit crowding agents that have an affinity towards other biomolecules. Our aim is to underscore the structural basis of accelerated protein transportation in nuclear environment by identifying the key molecular determinants associated with it. The study deals with bulk crowders and genomic crowders separately. The latter refers to the DBPs that are already bound to the DNA and spread over the entire genome covering ~20-50% of it(48). For example, 90% of LacI copies of a cell are hooked to DNA nonspecifically at various sites that are not the cognate site but shares a varying degree of sequence similarities with the target DNA site(49). These proteins serve as roadblocks to a searching protein during its 1D diffusion along the DNA. By performing extensive Langevin dynamics simulations, we show that the protein-crowder interactions play a pivotal role in enhancing the facilitated diffusion of DBPs inside crowded environment. Follow up structural analysis recognizes efficient target search modes of the searching protein actuated by crowder molecules. The proposed molecular picture is consistent with a cargo transferring mechanism, where crowder molecules with intermediate nonspecific crowder interactions transport the protein molecule quickly and effectively for faster recognition of the cognate DNA sites. The results are useful to study gene regulation kinetics of transcription factors and detailed models of facilitated diffusion in eukaryotes.

## MATERIALS AND METHODS

### Molecular systems

In this article, we used coarse-grained descriptions for protein, DNA and crowder molecules. Sap-1 (PDB ID: 1BC8)(50) is selected as a DBP that specifically binds with a 9-base pair DNA site of sequence ACTTCCGGT. We modelled the protein by replacing each amino acid with a sphere of 2 Å radius placed at the respective *C*_*α*_ positions (see Supplementary Fig S1). The protein maintains its folded structure during the simulation with the help of a native topology based Lennard-Jones potential(51) that promotes only the formation of contacts present in the experimentally determined folded structure of the protein. The DNA molecule is modelled by three beads per nucleotide, placed at the center of phosphate, sugar, and base respectively. The energy function is adopted from 3SPN.2C model of DNA developed by Pablo et al.(52). Further details regarding modelling of these two molecules are available in the supplementary text. The DNA model has been successful in predicting structural features of DNA such as major and minor grooves geometry as well as its mechanical properties including persistence length and melting temperature with respect to varying ion compositions(53, 54). We model the protein-DNA nonspecific interactions through short-range repulsive excluded volume interactions and long-range electrostatic interactions captured by Debye-Huckel potential. For electrostatic interactions, we considered unit positive charge on Arg and Lys amino acids and unit negative charge on Asp and Glu amino acids. For DNA, a net charge of −0.6 has been placed on every phosphate bead considering the effects of counter-ion condensation near the DNA surface. The application of Debye-Huckel potential has been widely studied in various nucleic acid and protein biophysics(55, 56) despite its limited applicability at low ion concentrations.

### Crowder model

The crowding agents are modelled as uncharged spheres that occupy a volume fraction, *ϕ* = 4*N*_*c*_ *π R*^3^/3*L*_*x*_*L*_*y*_*L*_*z*_, where *L*_*x*_, *L*_*y*_ and *L*_*z*_ represents the dimensions of the simulation box. *N*_*c*_ represents the total number of crowder molecules present in the system with a radius R set at 7.8Å. The value is consistent with the crowder dimension (PEG 600)(57) used in the experiments described in ref 27 that reported enhancement in the facilitated diffusion of glycosylase proteins in the presence of PEG crowding. Our objective is to investigate if the similar kind of crowding environment, but with nonspecific crowder affinity that PEG also features in contradiction to its believed ‘inert’ character, enhances the facilitated diffusion of DBPs, and if yes, to probe the underlying molecular mechanism. To this end, one should note that the present crowder model is not intended to mimic the other features of cytoplasmic crowders such as size heterogeneity that plays crucial roles in regulating various dynamic phenomena(41, 58, 59), rather to investigate how attractive interactions between searching protein and the crowder molecules affect the diffusion of the former. While a crowder model that mimics all the features of cytoplasmic crowders including their compressibility, malleability, size to charge ratio etc. is always desirable, it may not be straight forward to correlate its action directly with one of its molecular property.

In the present study, we consider the bulk and genomic crowders separately. For bulk crowders, we adopted *ϕ*=0.3 for all our simulation studies, which corresponds to the physiological cell crowding. The genomic crowders are modelled by unbiasedly placing the crowder spheres at the major grooves of the DNA at approximately equal distances. Their association with the DNA molecule is maintained through a pseudo bonded potential. The crowder molecules are allowed to interact with other crowders and protein, DNA through a pairwise potential is given by(60),

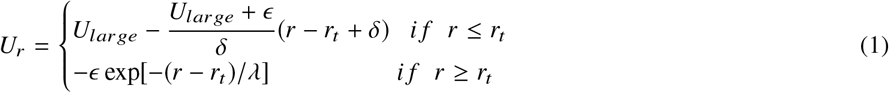

where *r*_*t*_ is the sum of the radii of crowders and interacting beads. The repulsive interaction is modelled by the first part of the equation whereas the second part is responsible for an attractive interaction. A large finite force, given by *U*_*large*/*δ*_ maintains the hard-core interaction approximation and *ϵ* (*k*_*B*_*T*) gives the attractive interaction strength acting within the characteristic range λ between crowders and other molecules. We set *U*_*large*_ at 40.0 kcal/mol and *δ*=1Å with a characteristic range =5 Å throughout our work. The attractive interaction is ignored beyond *r* − *r*_*t*_ > 6*λ*.

### Simulation Protocol

We study the motion of protein, DNA and crowder molecules through an overdamped Langevin dynamics simulation. The friction coefficient, γ is set to 0.05 at a temperature, T= 300 K and a physiological salt concentration of 120 mM. The simulations are performed by initially placing a 100-base pair (bp) B-DNA in the middle of a 150Å X 150Å X 410Å simulation box with periodic boundary condition. The protein is placed far from the DNA molecule, whereas the crowders are distributed randomly inside the simulation box. The initial DNA conformation is generated using w3DNA web server(61) containing Sap-1 binding site at the middle of a random DNA sequence. Upon reaching the target DNA site, the protein specifically binds with the DNA through a soft, attractive Lennard-Jones potential. Further details are given in Supplementary text. All our production runs are 1×10^8^ MD steps long during which the role of nonspecific crowder interactions on the DBP dynamics is monitored by varying the attractive interaction strength (*ϵ*) of the crowder molecules. For each *ϵ*, we performed 20 independent runs to investigate the detailed search mechanism with acceptable statistical significance.

We have also performed an all-atom simulation at 300 K to probe interactions between PEG and Sap-1. The simulation was performed using the GROMACS molecular dynamics package with the OPLS force field in the presence of PEG600 crowders. We used a base time step of 2 fs. The neighbour search was performed using the Verlet Algorithm and the Particle Mesh Ewald (PME) method that effectively treats the long range electrostatic interactions. We used the TIP3P water model to perform the simulations.

### Estimation of depletion layer dimension and characterizing Sliding, Hopping, Floating and 3D Diffusion dynamics of DBPs inside crowded milieu

We estimated the dimension (width, *l*_*d*_) of the depleted volume formed due to the presence of crowder molecules around DNA by measuring the average distance between the center of each DNA base pair and the nearest crowder molecule. *l*_*d*_ decreases when the protein approaches the DNA closer than twice the radius of the crowding agents. The newly formed depletion zone is referred as composite depletion zone that features a narrower width because of attractive depletion force. Schematic description of the formation of composite depletion zone is depicted in Fig S2.

To identify various search modes adopted by the diffusing protein, we follow the description similar to our previous works(40, 41). Briefly, the protein is said to perform sliding if at least 70% of the recognition helix of Sap-1 is within the DNA major groove with an orientation angle of < 25°, and the center of mass of the recognition helix lies within 8 Å to the closest DNA base pair in the absence of crowder molecules (see Fig S3). The associated electrostatic energy 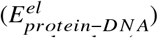 between protein and DNA ranges from −12 to −8 kcal/mole, indicating the strongest interaction between the two molecules (see Fig S4). The protein is assumed to be performing 3D diffusion if its recognition helix is more than 25 Å away from the closest DNA base pair. At this distance, protein-DNA electrostatics energy falls to less than 2 kcal/mol. Associated protein dynamics is assumed to be independent of DNA for such low interaction energy. In contrast, we consider the protein is performing hopping if it is not satisfying one or more criteria of sliding and is typically positioned 8-15 Å away from the closest DNA base pair. Corresponding strength of electrostatic interaction (−8 to −3 kcal/mol) ensures the protein to be close to the DNA molecule and not to get dissociated off the DNA surface completely. The protein, under such condition, performs small jumps (hopping) on the DNA. We consider the protein to be performing floating dynamics if it is 15Å −25Å away from the DNA surface and doesn’t satisfy any of the sliding criteria. The corresponding electrostatic energy (−3 to −1 kcal/mol) is inadequate to keep the protein close to the DNA surface that distinguish it from usual hopping dynamics, where protein jumps under the influence of DNA electrostatics, as explained above. During the floating dynamics, the protein is located inside the composite depletion zone and therefore experiences reduced viscosity than bulk solution. The 1D and 3D diffusion coefficients are estimated from the linear behaviour of the mean square displacement of the protein molecule during 1D (slide and hop) and 3D diffusion respectively.

## RESULTS AND DISCUSSION

In order to investigate how molecular crowding enhances DNA target search efficiency of DBPs, we study the diffusion of a transcription factor Sap-1 on a 100 bp linear DNA segment. We performed the experiments separately using explicit random crowder molecules (bulk crowders) and DNA bound crowders (genomic crowders). For both crowding agents, we systematically vary the strength of the nonspecific protein crowder interactions and probe the molecular picture of the target search mechanism of DBPs. These experiments help us to identify the molecular determinants responsible for faster search kinetics of DBPs inside crowded milieu.

### Bulk crowding modulates environment for 1D and 3D diffusion of DBPs

In the presence of bulk crowders, the protein moves mainly in two ways: either it diffuses three-dimensionally away from the DNA molecule (see Fig 1A) or one-dimensionally close to the DNA surface. During 3D diffusion, the protein interacts with the randomly moving crowder molecules and its overall diffusivity is governed by the concentration, size, shape, and mobility of the crowding agents(41). In comparison, protein diffusion during the 1D search regime was found to be significantly different in the presence of purely repulsive crowders(40, 41). This is due to the change in the search environment during 1D diffusion of the protein. Study indicates that when protein approaches the DNA closer than twice the radius of the crowder molecules (see Fig 1B), their individual depleted volumes merge due to an entropic force and the combined volume of the newly formed depletion zone around the protein-DNA complex (composite depletion zone(42)) decreases (see Fig S2). The micro-environment inside the depletion zone, which is spanned over a few nanometers over the DNA surface features much lower viscosity in the absence of crowder molecules compared to that of the bulk solution. The associated protein dynamics in this region is, therefore, independent of the crowder concentration and their physiological properties(41) and follows a 1D random walk along the DNA contour. Presumably, the effect would be different for interacting crowder molecules, where the enthalpic contribution from crowders needs to be taken into consideration while studying the formation of depletion zone around the DNA molecule and the protein diffusion inside it. The situation closely resembles to the nuclear environment, where proteins and other biomolecules frequently interact among themselves nonspecifically. We also enquire if the in vitro crowding agents such as poly ethylene glycol (PEG) interact similarly through nonspecific interactions. Towards this end, it should be noted that PEG that has been used extensively in the experimental studies and significant enhancement in the facilitated diffusion of DBPs was reported(27, 29). To investigate, we perform an all-atom MD simulation (see Materials and Methods for details) of Sap-1 with ~350mg/ml solution of PEG-600 crowders. The result in Fig 1C suggests preferential binding of PEG to Sap-1 as can be seen from the accumulation of PEG molecules (blue region) on specific surfaces of Sap-1 compared to the bulk solution. Few other studies along the line have confirmed similar active interactions of PEG molecules with biological milieu(62, 63). How such nonspecific crowder interactions affect the microenvironment of the target search dynamics of DBPs? We investigate this by simulating the protein diffusion on DNA in the presence of crowders, where nonspecific interaction strength of the crowders (*ϵ*/*k*_*B*_*T*) as varied gradually from 0 to 0.5. Our result in Fig 1D shows its direct impact on the width of the depletion region (*l*_*d*_), that decreases from 9.5 Å to 5.0 Å with the increasing *ϵ*. The range of *l*_*d*_ matches well with the experimentally observed depletion width of PEG-600(27). The decreasing trend in *l*_*d*_ is also accompanied by a simultaneous increase in protein-DNA interactions during the one-dimensional DNA search (increasing 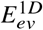), reflecting the rising protein-crowder crosstalk during this search regime.

**Figure 1:**
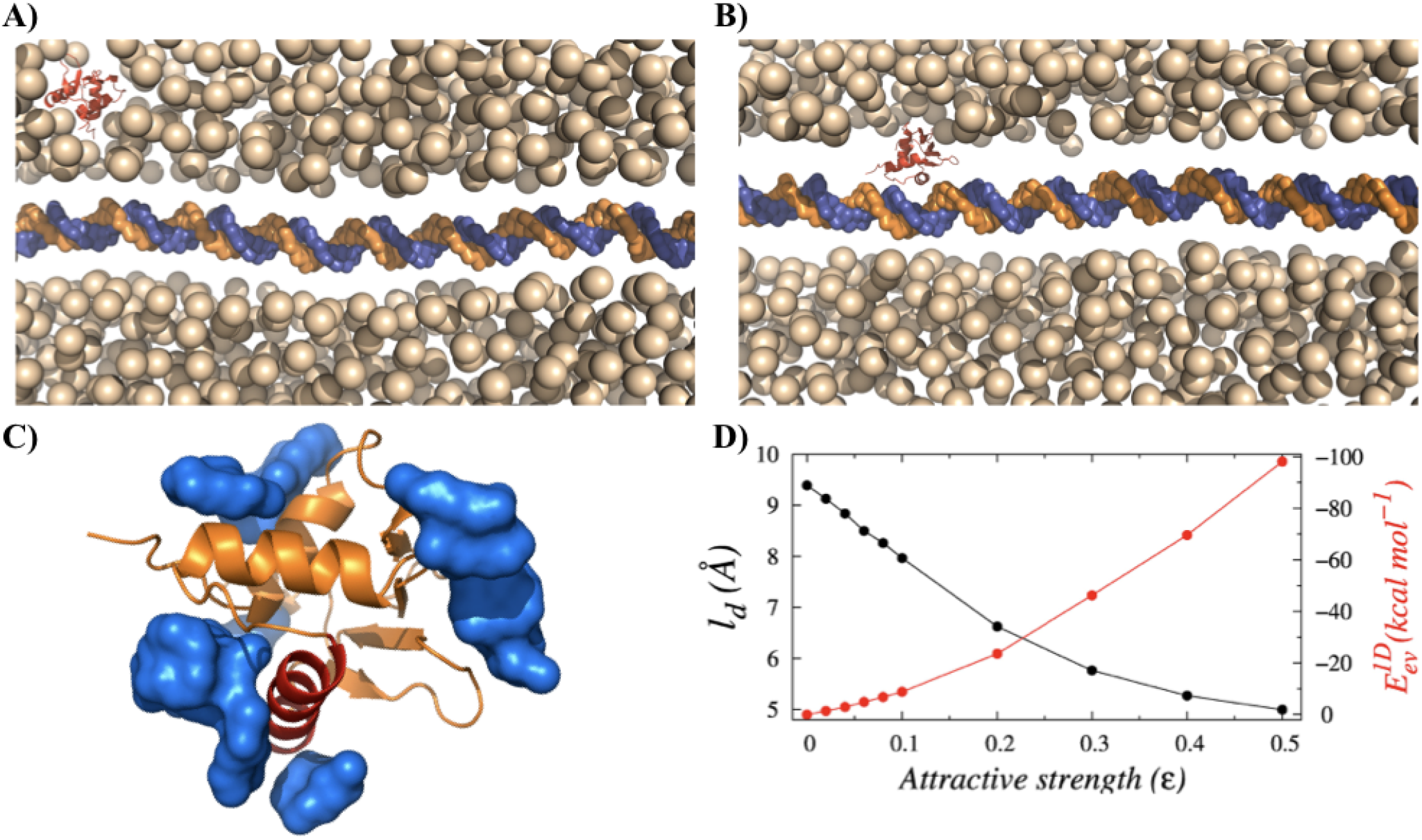
Schematic representation of the target search process of DNA Binding Proteins (DBPs) inside a crowded medium. (A) The crowding agents form depletion zones (white zone) around the protein and DNA molecule separately when they are far from each other (3D diffusion). (B) When the molecules are in close proximity, the depletion zones merge and the protein moves along the DNA contour one-dimensionally through the composite depletion zone. (C) During both 3D and 1D diffusions, the crowder molecules interact with the diffuser protein through nonspecific short-range attractive forces, as can be seen from the preferential association of PEG-600 crowders (shown in blue surface) on Sap-1 protein surface in an all-atom simulation. (D) Variations in width of the depletion region (width, *l*_*d*_) around the DNA molecule and excluded volume interactions 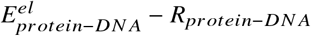 between the searching protein and the crowder molecules with the increasing attractive strength of crowders(*ϵ*).

### Protein diffusivity on DNA inside crowded milieu

How the synergistic actions of decreasing depletion width and increasing protein-crowder crosstalk with increasing strength of nonspecific protein-crowder interactions, *ϵ* influence the selection of protein search mode? To quantify this, we estimate the propensities of 3D and 1D search modes adopted by the protein to reach to the target DNA site and presented them as functions of *ϵ* in Fig. 2A. The results indicate that with increasing *ϵ*, the 1D propensity of the searching protein decreases whereas the 3D diffusion propensity increases. This can be rationalized from the fact that with increasing *ϵ*, the strong nonspecific attractive force from the crowder molecules pull the protein out of the composite depletion volume to the bulk where the protein translocation is 3D diffusion limited. The protein is positioned at least 25Å away from the DNA surface where it diffuses independent of the DNA electrostatics. In contrast, a small *ϵ* value ensures minimal interference from the crowding agents during the protein diffusion through the depletion zone under the influence of strong DNA electrostatics. This can be seen from the high sliding propensity of the protein at small *ϵ* value as presented in Fig S5. The strong DNA electrostatics keeps the protein at close vicinity and the protein diffuses along the helical pitch of the DNA slowly to read out the DNA bases. In Fig. 2B, we present the average displacement (*D*_*z*_) of the searching protein along the DNA contour during a 1D diffusion event, where a single event refers to a continuous period of time in a particular search mode that ends with the change in the search mode of the protein. The result suggests that with increasing strength of nonspecific protein-crowder interactions, diffusion length (*dz*_(1*D*)_) exhibits an initial rise up to *ϵ*=0.2 and then a rapid decrease. The trend indicates a trade-off between the electrostatic interactions from the DNA and nonspecific attractions from the crowding agents in the intermediate *ϵ* range that determine the non-monotonicity of the (*dz*_(1*D*)_) with varying *ϵ*. Similar behaviour can be observed for the average duration of hopping events as well as shown in Fig S6. How does such non-monotonic dependence of (*dz*_(1*D*)_) on the *ϵ* influence the overall target search efficiency of the protein?

**Figure 2:**
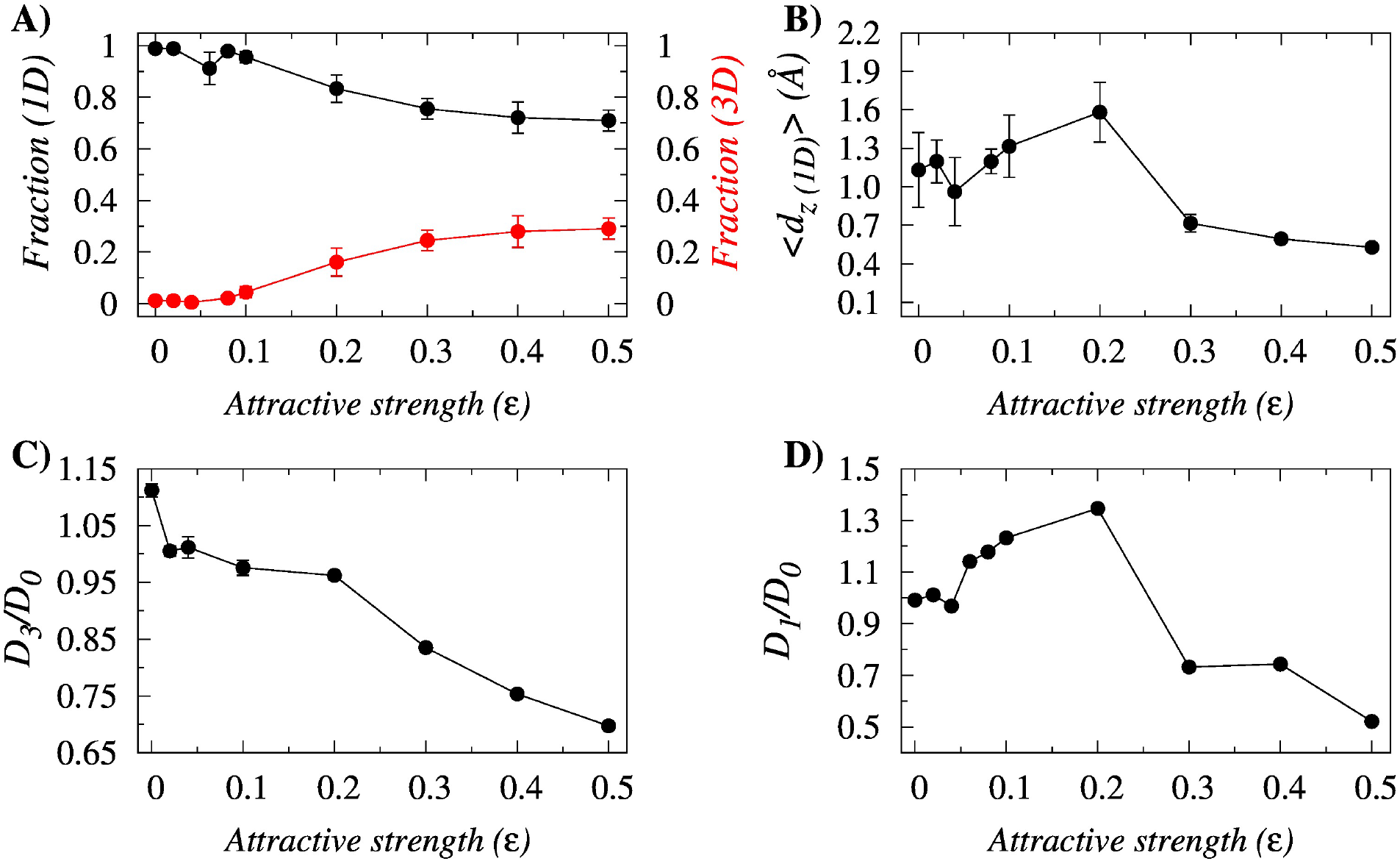
Effects of nonspecific crowder interactions (*ϵ*) on the target search mode of DBPs. (A) The variation in the affinities of 1D and 3D search modes of the searching protein as a function of the *ϵ*. (B) Variation in the average Z-displacement traversed by the Sap-1 protein per 1D event on the DNA surface as a function of *ϵ*. Impact of nonspecific crowder interactions on the diffusivity of the searching protein. (C) The variation in the relative 3D and (D) 1D diffusion coefficient of Sap-1 as functions of *ϵ*. In Fig. 2D, the associated error bars are smaller than point size. *D*_0_ represents the diffusion coefficients in the respective modes in the absence of crowder molecules.

To probe the issue, we separately estimate the 3D and 1D diffusion coefficients of the protein and present as a function of nonspecific crowder interactions (*ϵ*) with respect to the diffusivity measured in the absence of crowders and presented in Fig. 2C and 2D. Our result suggests first a slow and then a sharp decrease in the relative 3D diffusivity of Sap-1 with increasing *ϵ*. This is due to the growing stickiness of the crowder molecules with rising *ϵ* that holds the protein through strong nonspecific attractive forces and thereby hinders its free 3D diffusion. In contrast, the 1D diffusion coefficient of Sap-1 exhibits a nonmontonic dependence on *ϵ*, featuring a maximum at an intermediate *ϵ*=0.2 as shown in Fig 2D, suggesting the plausible role of longer 1D events at this *ϵ*. The corresponding acceleration in diffusion is 35% greater with respect to that in the absence of crowder molecules. To further realize how the protein molecule speeds up the search process during 1D dynamics and the role of crowders in it, we monitor the movement of the crowder molecules simultaneously with the diffusing protein. In Fig. 3A-3C, we present the time evolution of all crowder molecules at three different strengths of crowder affinities (*ϵ*). The fluctuating red lines indicate the indices of the crowder molecules that are interacting attractively with the diffuser protein at any given time. The evolving nature of the lines, therefore, indicates the footstep of the searching proteins monitored through the lens of interacting crowder indices. Our result shows that the unique indices of such interacting crowders decrease with increasing *ϵ* as presented in Fig 3D. Simultaneously, the transfer rate (*R*_*t*_) of the protein molecule that depends on the average time spent with the nearest crowder molecule increases. These signify that at low *ϵ*, the protein interacts weakly and nonspecifically with the crowding agents and therefore, is able to interact with many crowders while diffusing independently. In contrast, at high *ϵ*, the strong protein-crowder interactions allow the former to interact only with the adjacent crowder molecules (the number of unique crowder indices decreases in Fig 3D) but only for a short span of time. The strong pull experienced by the protein molecule from the surrounding crowder molecules forces it to shuttle among them that leads to a high transfer rate *R*_*t*_. The protein dynamics at this condition is strongly coupled with the dynamics of neighboring crowder molecules. At intermediate *ϵ*, the protein molecule adopts a combination of both independent as well as crowder regulated search dynamics that promotes longer hopping events and subsequently results in faster protein diffusion.

**Figure 3:**
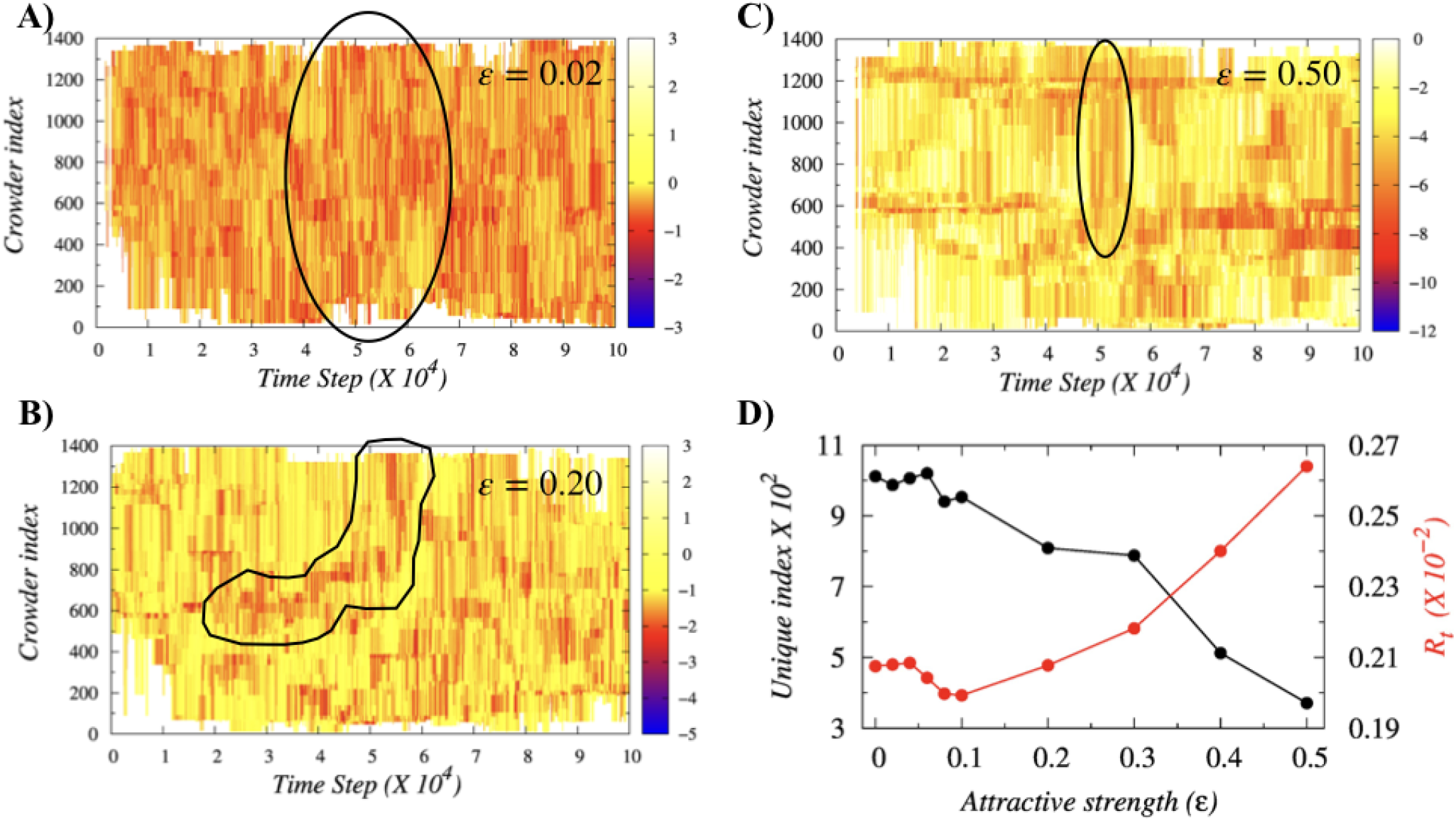
Protein diffusion through the lens of crowder dynamics. (A-C) Time evolution of indices of crowder molecules that are interacting with the searching protein through the nonspecific crowder interaction strengths *ϵ* = 0.02, 0.2 and 0.5 respectively. (D) The overall variations in the number of unique interacting crowder molecules and the rate of transfer *R*_*t*_ of Sap-1 (red line) from one crowder to another as a function of *ϵ*.

### Crowder induced search mode for facilitated diffusion

In order to capture the mechanistic details of the respective search dynamics, we monitor the time evolution of the distance (*R*_*protein-DNA*_) between the center of recognition helix of Sap-1 and center of the closest DNA base pairs in the presence of crowding agents of varying crowder affinities. The result in Fig 4A for *ϵ*=0.0 (purely repulsive crowders) shows that the protein performs primarily 1D diffusion along the DNA. Further inspection suggests that for *R*_*protein-DNA*_ < 8 Å, the protein is under strong influence of DNA electrostatics that orients the former such that the protein reads out the DNA base pair thoroughly in the sliding mode while rotating along the DNA helical pitch with simultaneous advancements along the DNA contour (rotation-coupled-sliding, see Fig S7). In comparison, when the protein is located at 8 Å < *R*_*protein-DNA*_ < 15 Å, it performs small jumps iteratively on the DNA surface (see movieS1). The DNA electrostatic interactions is moderate (−3 to −1 kcal/mol) at this condition that allows the protein to dissociate from the DNA surface for short time before it comes back again. One should also note here that a comparison of the associated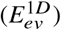 contour plots in the absence (Fig S4) and presence of entropic crowders (Fig 4B) suggests reduced probability of protein dissociation from the DNA surface (low population of protein for *R*_*protein-DNA*_ > 25Å) in the presence of crowding agents, indicating their role in promoting the 1D diffusion of DBPs.

**Figure 4:**
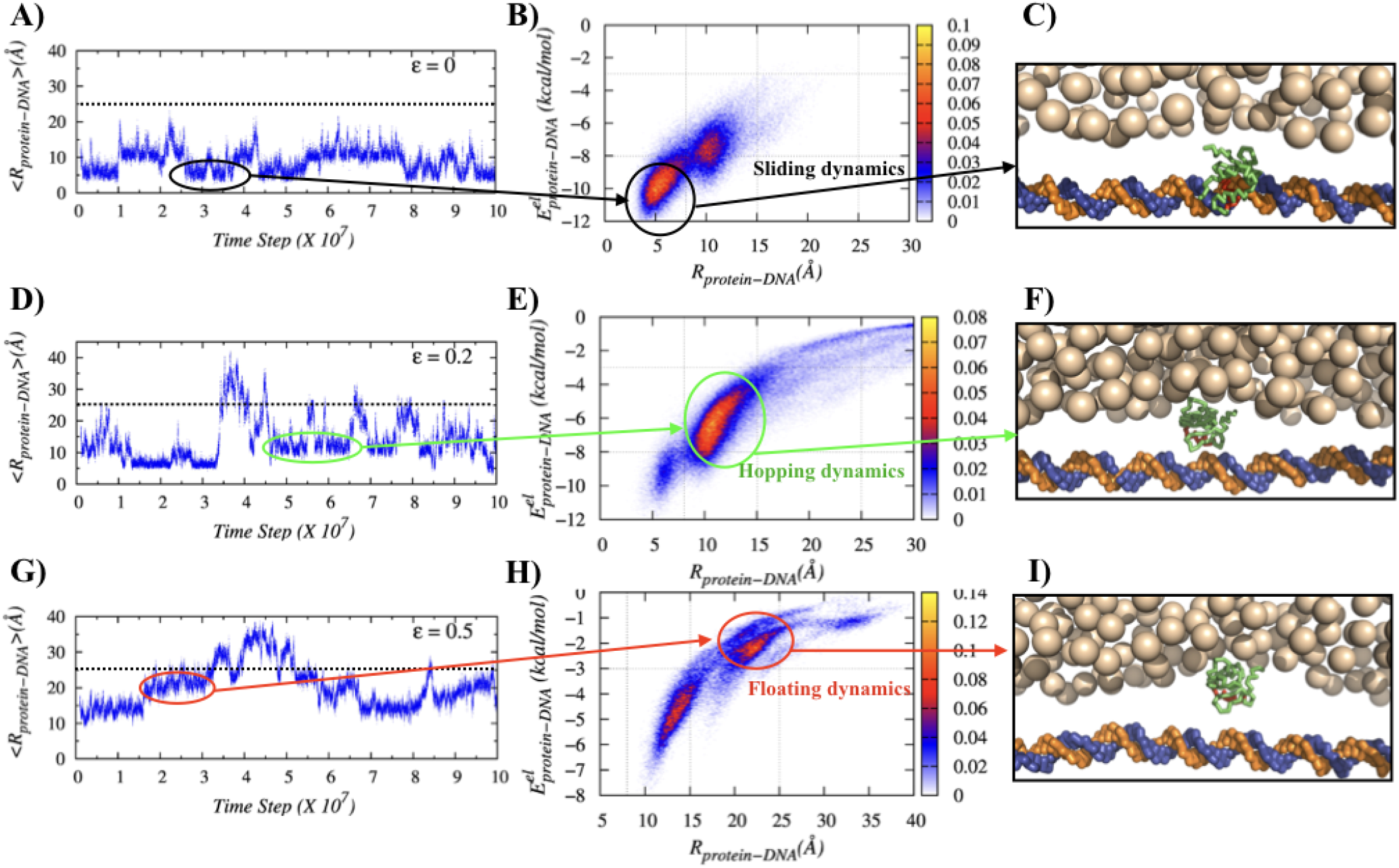
Structural basis of enhanced facilitated diffusion in presence of crowding agents. Horizontally the figures portray (i) the time evolution of distances between the center of recognition helix of Sap-1 and the center of the closest DNA base pair, (ii) 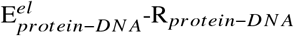 contour plots indicating the differences in various search modes and (iii) their corresponding snapshots showing the protein position during these search modes for nonspecific crowder interaction strengths,*ϵ* =0.0 (A-C), 0.2 (D-F) and 0.5 (G-I) respectively

For intermediate to strong nonspecific crowder interaction strengths, the sliding propensity diminishes (see Fig. S5 as well for propensities of various search modes with varying *ϵ*) as the protein-crowder nonspecific attraction is strong enough to pull the former off the DNA surface and promote hopping and 3D diffusion. Interestingly, here we find the protein to spend some times in a search mode at 15 Å 25Å away from the DNA bases. The associated protein-DNA electrostatic energy is inadequate (−3 to −1 kcal/mol, see Fig 4E and H) to make the protein hop. How the protein moves then along the DNA? We find that the protein-crowder interaction is the driving force. The crowder molecules transport the protein from one crowder to another (see movieS2) and thereby helps the protein to float on the DNA through the crowder free depletion zone. Estimation of the protein diffusivity (D1 coefficients, see Fig 5A) suggests that such crowder induced floating dynamics of the protein is ~36% - ~41% faster compared to that of the hopping dynamics. We also measure the impact of the floating dynamics on the overall search dynamics of the protein and find ~39% increment in the overall diffusion coefficient (*D*_0_) at *ϵ*=0.2 in Fig 5B compared to the protein diffusion in the absence of crowder molecules. The underlying molecular picture is revealed from the measurement of ruggedness (*σ*_*μ*_) of associated potential energy landscape following the prescription of Putzel et al(60) (for more details, see Supplementary text). Fig 5B suggests that *σ*_*μ*_ is minimum at moderate nonspecific crowder interactions. This is because of the weak DNA electrostatic field experienced by the searching protein and the absence of crowder molecules inside the depletion zone when the protein is positioned at *R*_*protein-DNA*_ 15Å-25 Å. A lower or higher *ϵ* either favours sliding or 3D diffusion during which the protein experiences highly rugged energy landscape originated from either strong protein-DNA electrostatic interactions or intense nonspecific protein-crowder interactions respectively. The effective nonspecific crowder interactions towards the diffuser protein can also be increased by simply enhancing the concentration of crowding agents of a given *ϵ*. Following the argument, we find in Fig S8 that protein diffusion first rises with increasing crowder concentration(*ϕ*), peaks at an intermediate *ϵ* value and then decreases sharply for a fixed *ϵ*=0.2. The result is in line with the experimental observation by Tan et. al.(26), who have reported non-monotonic dependence of gene expression rate on the concentration of PEG 8k crowders, featuring an initial increase, then decrease and finally a complete halt with rising crowder concentration.

**Figure 5:**
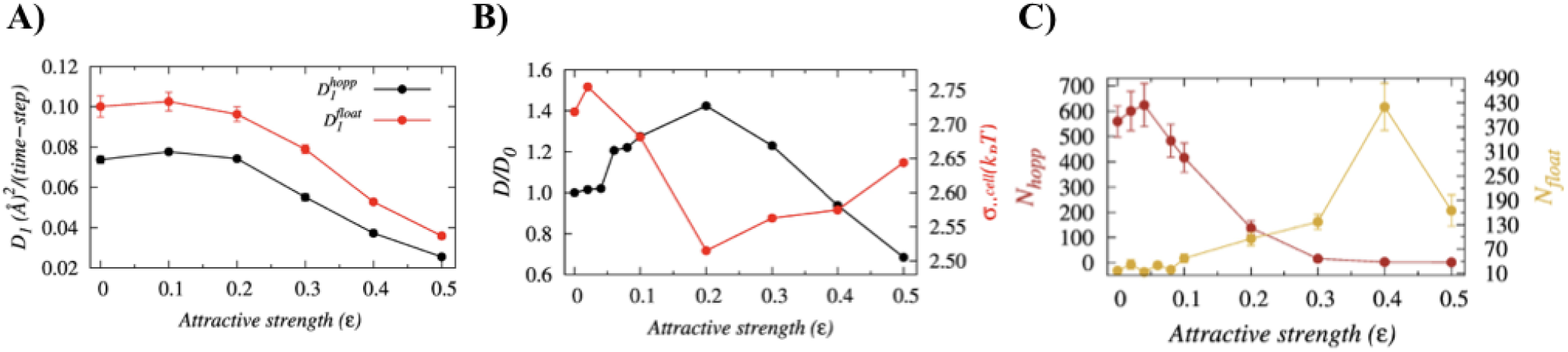
Impact of crowder induced floating dynamics of Sap-1 on the overall target search process. (A) Comparison between the efficiencies of hopping and floating dynamics as a function of crowder affinity (*ϵ*). (B) The variation in the overall diffusion coefficient (black like) of the protein (1D+3D, black lines) normalized by its value in the absence of crowders. The red line suggests the ruggedness of the associated potential energy landscape (red lines) on which the protein diffuses.(C)Variation in the total number of floating (*N*_*float*_) and hopping (*N*_*hop*_) events as a function of *ϵ*.

To this end, it may be noted that often genomic DNA contains innumerable sites that share high sequence similarities with the target DNA site. These pseudo cognate sites work as an antenna to pull down the searching protein onto the DNA surface through attractive interactions and thereby hinder the protein diffusion(64). It is tempting to speculate that the proposed floating dynamics induced by the bulk crowder molecules with moderate nonspecific crowder interactions could be a way to bypass the antenna-effect in order to expedite the search process. In other words, the proteins and other biomolecules present in the nuclear environment that can moderately interact nonspecifically with the searching DBPs may help the latter in bypassing pseudo cognate sites. We however, emphasise that the floating dynamics alone cannot assure the fastest target search kinetics, rather it requires an optimal balance between both the hopping and the floating dynamics. Our analysis at *ϵ*=0.2 suggests the balance is achieved for when the number of hopping (*N*_*hop*_) events is ~1.5 times to that of floating (*N*_*float*_) events (see Fig 5C).

### Facilitated diffusion in the presence of genomic crowders

Having seen the role of nonspecific crowder interactions in accelerating the facilitated diffusion of DBPs, we now turn to investigate the influence of interacting genomic crowders that are abundant in nature (~50% of the DNA surface is occupied by a vast majority of different kinds of proteins)(48). It has been noted that often such genomic crowders act as roadblocks to nearby searching proteins and hinder their diffusion(65, 66). In contrast, few other studies have reported marginal to significant rise in diffusion efficiency of DBPs in the presence of genomic crowders(33, 67). Clearly, the prevailing ambiguity requires deeper insights, particularly when the crowders may nonspecifically interact with the diffuser protein. Does the interaction retard diffusion of nearby proteins by halting them as many pseudo cognate sites do? We investigated the issue by estimating the 1D diffusion coefficient of the protein as a function of nonspecific crowder interactions (*ϵ*) of the DNA bound crowders. Fig 6A presents a schematic representation of the system and Fig 6B suggests a non-monotonic dependence of the 1D diffusion coefficient on nonspecific crowder interactions, *ϵ* with a maximum observed at *ϵ*=0.06. The associated D1 enhances by ~12% compared to when there is no genomic crowders present. The measurement of the potential energy ruggedness confirms the least ruggedness at *ϵ*=0.06 on which the protein can quickly diffuse along the DNA contour. The same can be realized from the contour plots as well, presented in Fig 6C-E. The spots on the plot show the position of the protein during our simulations. Interestingly, the protein positions in all these plots are not exactly aligned with the position of genomic crowders, rather they are more probable in between two consecutive genomic crowders. How is it connected with protein diffusion?

**Figure 6:**
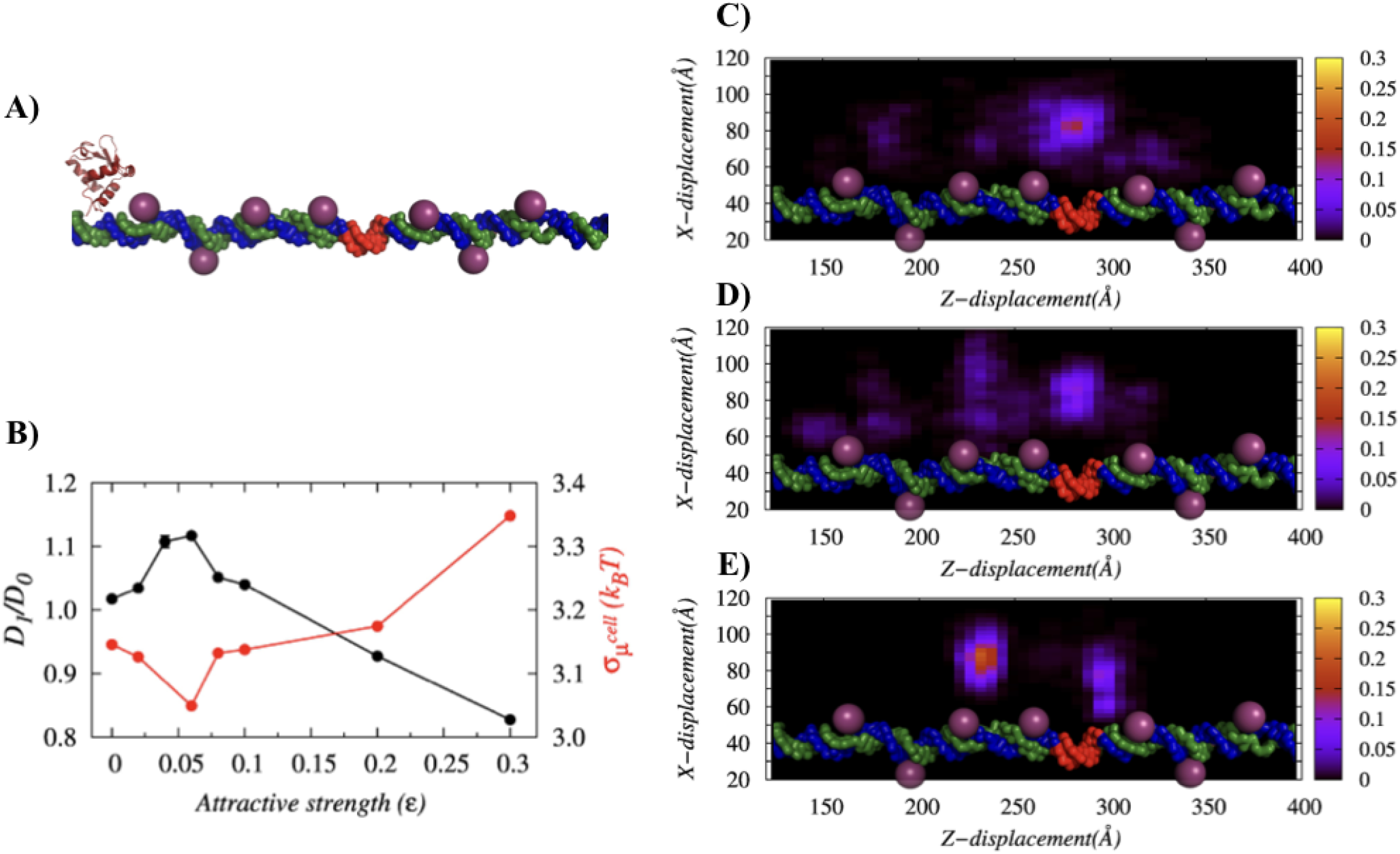
Role of genomic crowders on the target search process of DBPs. (A) Schematic representation of the target search process of Sap-1 (in orange) in the presence of DBPs that are already bound to the DNA (presented through spheres). (B) The variation in the 1D diffusion coefficient (in black lines) of the protein normalized by the diffusion coefficient in the absence of crowder molecules (*D*_0_) and the variation in the ruggedness of the potential energy landscape (in red lines) as a function of *ϵ*. (C-E) Contour plots showing the probability of protein positions on the X-Z plane for nonspecific crowder interaction strengths, (*ϵ*=0, *ϵ*=0.06 and *ϵ*=0.5 respectively. Brighter the spot, higher is the probability of finding the protein at that position.

A detailed structural analysis suggests that nonspecific crowder interactions allow adjacent or spatially close genomic crowders to interact with the nearby searching protein simultaneously, through the formation of a transient bridged complex (see Fig 7B). For genomic crowders on linear DNA, such interactions require deformation in the DNA conformation to bring the interacting crowders spatially close. In Fig 7A, we measure the DNA deformation through the angle (*θ*) formed between two adjacent genomic crowders and present as a function of nonspecific protein crowder interaction strength. The results imply that for moderate ϵ, the kink angle *θ* is minimum to promote the jumping of the protein from one crowder to another genomic crowder through the local deformation in DNA conformation and subsequently formation of a transient bridged complex. For purely repulsive crowders, formation of such bridged complex is not permitted. Similarly, the strongly attractive genomic crowders (high ϵ) hold the searching protein tightly and thereby unlikely to share it with other crowders for the formation of a bridged complex. The trend is reflected from the reduction in associated deformation in DNA (*θ*). To this end, we should emphasize the fact that the interaction strength of cytoplasmic crowders can be very weak or significantly strong to offset the facilitated diffusion through the transient bridged complex, which is observed for moderately interacting crowders. This may hinder the target search dynamics of the proteins. Other factors such as positions, mobility and size of genomic crowders also play crucial role in modulating the target search efficiency of DBPs(65, 66, 68). Nevertheless, specific DBPs have alternative mechanism to bypass the genomic crowders(69) by promoting formation of DNA loops and coils that are known to facilitate the target search process of proteins(70, 71).

**Figure 7:**
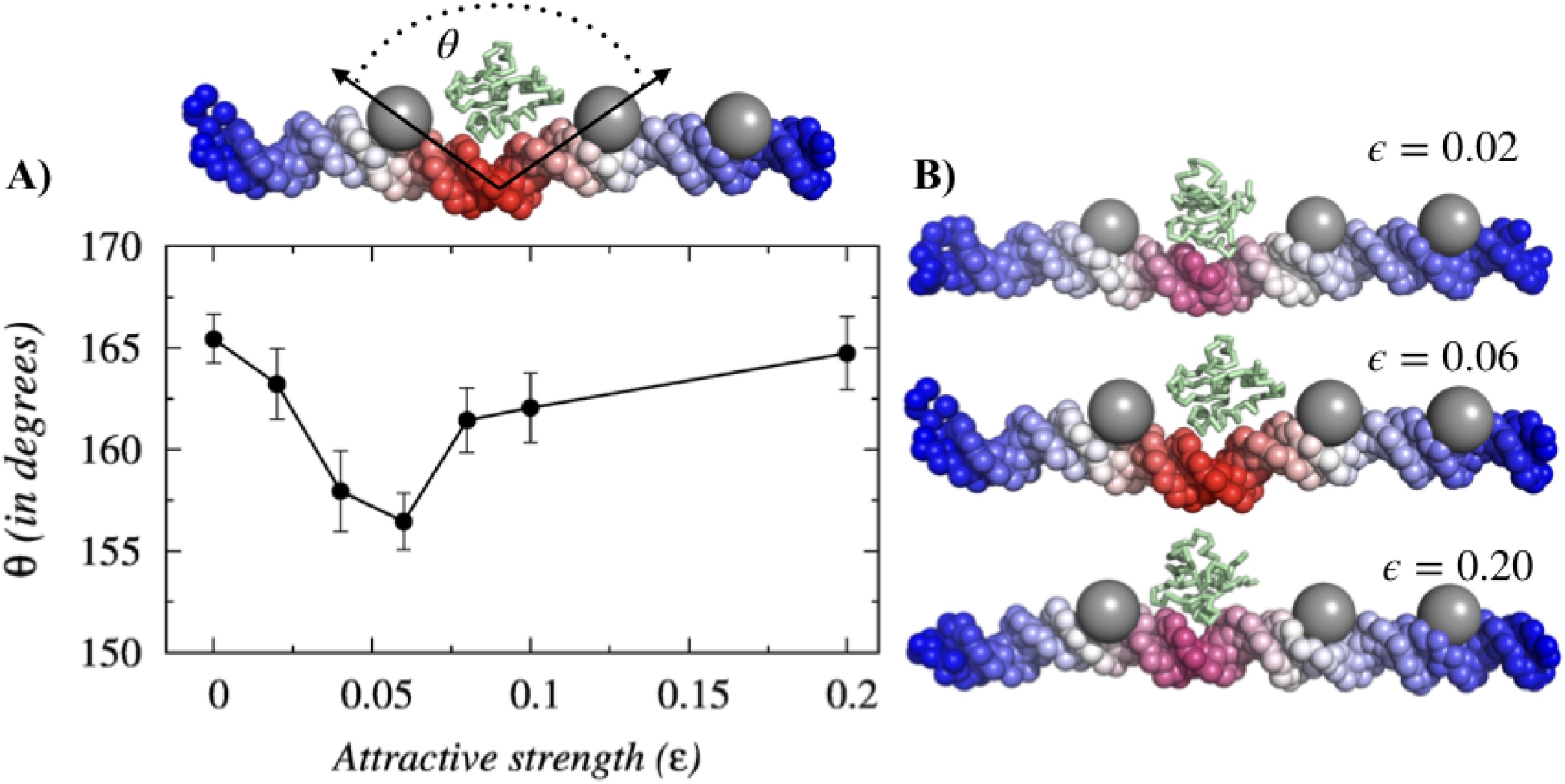
Crowder induced kinks in the DNA conformation and subsequent formation of a transient bridged complex. (A) Variation in the degree of the DNA deformation *θ* as a function of *ϵ*. The deformation originates when Sap-1 interacts with adjacent genomic crowders attractively. The crowder molecules tend to approach closer to the interacting protein and thereby causing local kinks in the DNA structure. (B-D) Schematic representation of the kink formed in the DNA structure for three different nonspecific crowder interaction strengths, (*ϵ*=0.02, *ϵ*=0.06 and *ϵ*=0.2). The increasing degree of bending is represented by the change in colour from blue to red.

## CONCLUSION

Through extensive molecular simulations of a coarse-grained model, we investigated the molecular mechanism of how the facilitated diffusion of DBPs enhances in the presence of crowded environment, where the crowding agents are capable of interacting with other biomolecules through attractive forces as in vivo crowders do. The model is successful in capturing the formation of depletion zone around a DNA molecule with a dimension comparable to that observed in the presence of a synthetic crowder (PEG 600). Any change in the crowder interaction strength is found to have a direct impact on the dimension of the depletion zone. The analysis, therefore, finds the crowder interaction strength as a key component in investigating the DNA target search dynamics of DBPs in a crowded medium. For example, we observe that the crowder molecules with moderate interaction strength expedite the search kinetics of DBPs maximally compared to when no crowder molecules are present. The counterintuitive result is understood from the structural analysis of simulation trajectories that identifies kinetically efficient alternative search modes of DBPs induced by the crowding agents. For bulk crowders, the alternative search mode resembles a protein floating one-dimensionally along the DNA contour. Unlike to the hopping dynamics during which protein iteratively performs small jumps near the DNA surface under the influence of the electrostatic field of the latter, the protein in the floating dynamics diffuses significantly away from the DNA surface, where it feels the DNA electrostatic field only marginally. The tradeoff between weak DNA electrostatics and moderate crowder interaction present a potential energy landscape with minimum ruggedness, suitable for maximum gain in search kinetics. On a similar note, the genomic crowders with moderate ϵ enhance the facilitated diffusion of a nearby interacting protein by transporting it from one to another adjacent crowder through the formation of a transient bridged complex between protein and adjacent genomic crowders. The DNA conformation subsequently adopts a locally kinked conformation to facilitate the process and helps the protein to bypass scanning the intermediate DNA bases in order to achieve faster kinetics. Both the proposed search modes are viable to only intermediate nonspecific crowder interactions. A stronger or weaker crowder interactions either holds back the protein for long and retard the search process or only marginally influences the protein dynamics. The emerged molecular picture is also consistent with the nonmonotonic dependence of the target search efficiency of DBPs on bulk crowder density observed by Tan et al.(26) and is useful to study gene regulation kinetics of transcription factors and facilitated diffusion in crowded cellular environment.

## AUTHOR CONTRIBUTIONS

A.B. and P.D. designed the research; P.D., and A.B. performed the research; P.D., and A.B. analyzed the data; P.D and A.B. wrote the paper.

## ACKNOWLEDGMENTS

We gratefully acknowledge the financial support from DST India (DST/INSPIRE/04/2013/000100, DST-SERB ECR/2016/000188, DST PURSE) and JNU (UPoE) research grants.

## SUPPLEMENTARY MATERIAL

Supplementary Material is attached

