## Supplementary for "Structural Basis of Enhanced Facilitated Diffusion of DNA Binding Proteins in Crowded Cellular Milieu"

P. Dey and A. Bhattacherjee

**List of contents:** **Page no.**

1. Simulation models 2-6
    a. Protein model. 2-3
    b. DNA model. 4-6
    c. Crowder model. 6-7
    d. Protein-DNA interactions. 7
2. System Configuration and Preparation 8
3. Criteria for data analysis 8-9
4. Tables 10-11
5. Figures 12-17
6. Supporting References 17-18

### 1. Simulation models.

**a. Protein model:** The protein used for our study is as a 93 residue long transcription factor, Sap-1(PDB ID: 1BC8) that binds to a 9-bp DNA target site(1). The single domain protein contains a well-defined recognition helix (53 to 68 residues, see the red region in S1) to scan and make specific contacts with the target DNA site. It is modelled by representing each amino acid as one bead placed at the respective Cα positions. The folded structure of the protein is ensured by a structure based Leonard-Jones potential(2) throughout the simulation. Moreover, a Debye-Huckel potential models the electrostatic interactions between the negatively charged amino acids (Glu and Asp) and the positively charged amino acids (Arg and Lys). Despite the limited use of Debye-Huckel potential to dilute ion solutions, it has successfully predicted many crucial aspects of nucleic acid biophysics(3–6).

The potential used to model the protein molecule is given as,

$$E_{pot} = E_{bond} + E_{bend} + E_{torsion} + E_{LJ} + E_{ev} + E_{ele} \quad (1)$$

The bonded energy, $E_{bond}$ is given as,

$$E_{bond} = \sum_i k_b (r_i - r_i^0)^2 \quad (2)$$

where $k_b$ =23.90 kcal/mol/Å$^2$, $r_i$ and $r_i^o$ are the distances between $i$-th and $i$+1-th $C_\alpha$ beads in an intermediate and the folded structures of Sap-1 respectively.

The potential energy function for any variation in angles, $E_{bend}$ is given as,

$$E_{bend} = \sum_i k_\theta (\theta_i - \theta_i^0)^2 \quad (3)$$

where $k_\Theta$ = 4.78 kcal/mol/rad$^2$, $\Theta_i$ and $\Theta_i^0$ are the angles among $i$-th, $i$+1-th, $i$+2-th $C_\alpha$ beads in an intermediate and the folded structures of Sap-1respectively.

The potential energy function for torsional angle between four atoms connected by bonds, $E_{torsion}$ is given as,

$$E_{torsion} = \Sigma_i\{k_{\phi 1}[1 - cos3(\phi_i - \phi_i^0)] + k_{\phi 2}[1 - cos(\phi_i - \phi_i^0)]\} \quad (4)$$

where $k_{\phi 1}$=0.119 kcal/mol, $k_{\phi 2}$=0.239 kcal/mol, $\phi_i$ and $\phi_i^0$ denotes the torsional angles between *i*-th, *i*+1-th, *i*+2-th, *i*+3-th $C_\alpha$ beads in an intermediate and the folded structures of Sap-1 respectively.

$E_{LJ}$ estimates the conformational energy using a native topology based model(2) in which the formation of contacts found in the folded structure of the protein is modelled by a Lennard–Jones potential.

$$E_{LJ} = \Sigma_{i<j-3}^{native}\epsilon_{ij}\left[5\left(\frac{\sigma_{ij}}{r_{ij}}\right)^{12} - 6\left(\frac{\sigma_{ij}}{r_{ij}}\right)^{10}\right] \quad (5)$$

where $\epsilon_{ij}$=3.824091778 kcal/mol. $r_{ij}$ and $\sigma_{ij}$ are the distances between the native pairs in a snapshot generated at a given time and in the folded structure of the protein respectively.

$E_{ev}$ represents the excluded volume interactions estimated between all non-bonded and non-native pairs of Sap-1 and also between the protein and DNA and is given as,

$$E_{ev} = \Sigma_{i<j-3}^{non-native}\varepsilon_{ev}\left(\frac{\sigma_{ij}}{r_{ij}}\right)^{12} \quad (6)$$

where $\varepsilon_{ev}$=0.239 kcal/mol, $r_{ij}$ gives the distance between *i*-th and *j*-th beads and $\sigma_{ij} = \sigma_i + \sigma_j$ represents the interaction specific length scale, where $\sigma_i$ and $\sigma_j$ are the interacting bead's radius.

The electrostatic interactions is modelled by Debye-Hűckel potential, $E_{elec}$, and is given as,

$$E_{elec} = \sum_{i<j}\frac{q_i q_j e^{-r_{ij}/\lambda_D}}{4\pi\epsilon_0\epsilon(T,C)r_{ij}} \quad (7)$$

where $q_i$ and $q_j$ denotes the charges on site $i$ and $j$, $r_{ij}$ represents the separation between the two sites. $\epsilon(T,C)$ is the dielectric permittivity of solution and is a function of the molarity of NaCl and temperature[4] as

$\in(T,C) = \in(T)a(C)$ ,

$\in(T) = 249.4 - 0.788\,T/K + 7.20\times10^{-4}(T/K)^2$ and,

$a(C) = 1.00 - 2.551\ C/M + 5.151\times10^{-2}(C/M)^2 - 6.889\times10^{-3}(C/M)^3$

The Debye screening length is given as;

$$\lambda_D = \sqrt{\frac{\epsilon_o \epsilon(T,C)}{2\beta N_A e_c^2 I}}$$

β is the inverse thermal energy of the system $(k_B T)^{-1}$, $k_B$ denotes the Boltzmann constant, $N_A$ is Avogadro's number and $I$ is the ionic strength of solution.

**b. DNA model:** We describe the DNA force field by the 3SPN.2C model by Pablo et.al.(7) In this model, each nucleotide is represented by three beads placed at the geometric centre of phosphate, sugar and base respectively. This model accurately estimates the structural properties such as, major and minor grooves which is consistent with experimental results. The model is able to correctly capture the persistence length for both ss- and ds-DNA. Also, it can the predict dsDNA melting temperatures that are in good agreement with experimental results. These features make it a suitable candidate to study the molecular basis of DNA dynamics.

The total potential energy function of the DNA is given as,

$$E_{pot}^{DNA} = E_{bond}^{DNA} + E_{bend}^{DNA} + E_{tors}^{DNA} + E_{exe}^{DNA} + E_{bstk}^{DNA} + E_{cstk}^{DNA} + E_{bp}^{DNA} + E_{elec}^{DNA} \qquad (8)$$

The bonding energy, $E_{bond}^{DNA}$ is given as,

$$E_{bond}^{DNA} = \sum_i k_b (r_i - r_i^o)^2 + 100 k_b (r_i - r_i^o)^4 \qquad (9)$$

where $k_b$ = 0.6 kJ/mol/Å$^2$ and $r_i$, $r_i^0$ are the instantaneous and equilibrium bond length for $i$-th bond respectively.

The energy function for bending, $E_{bend}^{DNA}$ is given as,

$$E_{DNA}^{bend} = \sum_i k_\theta (\theta_i - \theta_i^0)^2 \tag{10}$$

Where $k_\Theta$ = 200 kJ/mol/rad$^2$, $\theta_i^0$ represents the instantaneous and equilibrium bond angles for i-th bond angle respectively.

$E_{tors}^{DNA}$ denotes the potential energy function for torsional angle between every four atoms connected by bonds,

$$E_{tors}^{DNA} = \sum_i - k_\phi \exp\left(\frac{-(\phi_i - \phi_i^0)^2}{2\sigma_{\phi,i}^2}\right) \tag{11}$$

where $k_\phi$ = 6.0 kJ/mol, the well-depth, equilibrium angle, and Gaussian well-width of dihedral *i* are given by $\phi_i$ , $\phi_i^o$ and $\sigma_{\phi,i}$ respectively.

$E_{exe}^{DNA}$ denotes the energy function for excluded volume between sites *i* and *j* and is given as,

$$E_{exe}^{DNA} = \sum_{i<j} \begin{cases} \epsilon_r \left[\left(\frac{\sigma_{ij}}{r_{ij}}\right)^{12} - 2\left(\frac{\sigma_{ij}}{r_{ij}}\right)^{6}\right] + \varepsilon_r & , r < r_c \\ 0 & r \geq r_c \end{cases} \tag{12}$$

It is a purely repulsive potential where $\epsilon_r$=1.0 kJ/mol denotes the energy parameter, $\sigma_{ij}$ is the average site diameter between *i* and *j*, and $r_{ij}$ denotes the separation between them and $r_c$ is the cutoff distance for the excluded volume interactions.

The potential energy for intra-stand Base-stacking, $E_{bstk}^{DNA}$ is given as,

$$E_{bstk}^{DNA} = \sum_{bstk} \begin{cases} U_m^{rep}(\epsilon_{ij}, \alpha_{BS}, r_{ij}) + (K_{BS}, \Delta\theta_{BSij}) U_m^{attr}(\epsilon_{ij}, \alpha_{BS}, r_{ij}) & r_{ij} < r_{ij}^0 \\ f(K_{BS}, \Delta\theta_{BSij}) U_m^{attr}(\epsilon_{ij,} \alpha_{BS}, r_{ij}) & r_{ij} \geq r_{ij}^0 \end{cases} \tag{13}$$

where $K_{BS}$ = 6.0, $\alpha_{BS}$=3.0 and $\epsilon_{ij}$ represents the depth of the well of attraction between sites i and j, the equilibrium separation between the sites is given by $r_{ij}^0$, and $\alpha_{BS}$ serves to adjust the range of attraction. The decomposition of attractive and repulsive portions of Morse potential are modulated by the function, *f* using $\theta_{BS}$ .

The potential energy function for base pairing, $E_{bp}^{DNA}$ is, (14)

$$E_{bp}^{DNA} = \sum_{bp} \begin{cases} U_m^{rep}(\epsilon_{ij}, \alpha_{BP}, r_{ij}) + \frac{1}{2}(1+\cos(\Delta\phi_i)) f(K_{BP}, \Delta\theta_{1ij}) f(K_{BP}, \Delta\theta_{2ij}) U_m^{attr}(\epsilon_{ij}, \alpha_{BP}, r_{ij}) & r_{ij} < r_{ij}^0 \\ \frac{1}{2}(1+\cos(\Delta\phi_i)) f(K_{BP}, \Delta\theta_{1ij}) f(K_{BP}, \Delta\theta_{2ij}) U_m^{attr}(\epsilon_{ij}, \alpha_{BP}, r_{ij}) & r_{ij} \geq r_{ij}^0 \end{cases}$$

where $K_{BP}$=12.0, $\alpha_{BP}$=2.0 and $\epsilon_{ij}$ is the depth of the well of attraction between i and j, $r_{ij}^0$ represents the equilibrium separation between the two sites, and the parameter to control the range of attraction is given by $\alpha_{BP}$. *f* modulates the decomposition of attractive and repulsive portions of the Morse potential, The deviations from a reference dihedral angle is penalized by $\Delta\phi_1=\phi_1-\phi_1^0$. The decomposition maintains the repulsive character. *f* modulates the base pairing interactions using $\theta_1$ and $\theta_2$.

The potential energy function for cross stacking, $E_{cstk}^{DNA}$ is given as,

$$E_{cstk}^{DNA} = \sum_{cstk} f(K_{BP}, \Delta\theta_{3ij}) f(K_{cs}, \Delta\theta_{CSij}) U_m^{attr}(\epsilon_{ij}, \alpha_{CS}, r_{ij}) \quad (15)$$

where $K_{CS}$= 8.0, $\alpha_{CS}$= 4.0 and $\epsilon_{ij}$ gives the depth of well of attraction between sites i and j. $\theta_3$ and $\theta_{CS}$ modulates the cross stacking interactions.

$E_{elec}^{DNA}$ is a screened electrostatic potential acting between intra-strand and inter-strand phosphates and is modelled using the potential energy function given in Eqn. (7).

**c. Crowder model:** Crowders are modelled as uncharged spheres occupying a volume fraction, $\phi = 4N_c\pi R^3/3\, L_x L_y L_z$. Here, $L_x$, $L_y$ and $L_z$ are the dimensions of simulation box used with a periodic boundary condition, $N_c$ denotes total number of crowders and R is the radius of each crowder. Initially, the crowders are randomly placed inside the simulation box. Here, we describe the crowders that are present in the solution as bulk crowders and crowders occupying positions on the DNA surface as roadblocks. For bulk crowders, we adopted $\Phi$ =0.3 for all our simulation studies. The roadblocks are static and distributed uniformly on the DNA surface. The concentration of crowder molecules used in our study lies in the dilute regime to avoid several complex phenomena such as phi condensation etc. In our model, both the bulk crowders and roadblocks interact by a potential that includes both attractive and repulsive terms. The form of the pairwise potential is given by(8)

$$U_r = \begin{cases} U_{large} - \frac{U_{large}+\epsilon}{\delta}(r - r_t + \delta) & \text{if } r \leq r_t \\ -\epsilon \exp[-(r - r_t)/\lambda] & \text{if } r \geq r_t \end{cases} \tag{16}$$

where $r_t$ is the sum of the hardcore radii of crowders and interacting beads. The repulsive interaction is modelled by the first part of the equation whereas the second part gives rise to an attractive interaction. A large finite force, given by $U_{large}/\delta$ maintains the hard core interaction approximation and $\epsilon$ ($k_B T$) gives the attractive interaction strength acting within the characteristic range λ of crowders interacting with other molecules. Here, $U_{large}$ is set at 40.0 kcal/mol and δ=1Å with a characteristic range λ=5 Å. The attractive interaction is ignored beyond $r - r_t > 6\lambda$.

The sizes and masses of macromolecular crowding agents used in our study correspond to PEG600 crowders(9) used in experimental studies.

**d. Protein-DNA interaction:** In our model, protein and DNA are allowed to interact through two modes of interactions:

**(i) Non-specific interactions:** The Sap-1 protein non-specifically scans the DNA in quest of the target DNA site. During this process, two main interactions come into play. One is the electrostatic interaction between the charged amino acids and the phosphates present in DNA. Another nonspecific interaction is the excluded volume interaction between them. The electrostatic interaction is modelled using Debye-Huckel potential (Eqn.7) and excluded volume interactions are modelled using Eqn.6.

**(ii) Specific protein-DNA interactions:** When the protein reaches the target site, it binds specifically to the major groove using its recognition helix. The required information regarding the specific contacts is obtained from the X-ray crystallographic structure of Sap-1 (PDB ID: 1BC8). The formation of specific contacts are modelled using a short-ranged Lennard-

Jones potential as,

$$E_{LJ} = \Sigma_{i<j} \begin{cases} k_{sp}\left[5\left(\frac{\sigma_{ij}}{r_{ij}}\right)^{12} - 6\left(\frac{\sigma_{ij}}{r_{ij}}\right)^{10}\right], r < r_c \\ 0 \qquad\qquad\qquad\qquad\qquad\quad , r \geq r_c \end{cases} \tag{17}$$

where $k_{sp}$=0.5 kcal/mol, $\sigma_{ij}$ denotes the average site diameter between sites $i$ and $j$, while $r_{ij}$ is the separation between them. $r_C$ denotes the cut off distance at ($\sigma_{ij}$+5) Å.

### 2. System configuration and Preparation:

The coarse-grained model of the protein is generated from the atomic coordinates of the crystal structure, 1bc8.pdb. The 100 bp B-DNA structure, obtained from w3DNA (3D DNA structure) web server (http://w3dna.rutgers.edu)(10), provides the template for generation of coarse-grained model of DNA. The DNA is placed in the middle of the simulation box with a box size of 150Å X 150Å X 410Å. The formation of specific contacts are ensured by inserting a 9 bp target DNA sequence as found in the crystal structure of Sap-1 in the present DNA model. Initially, the protein is placed far from the DNA surface located at the centre of the simulation box along the Z axis and the crowders are distributed randomly inside the simulation box. We studied the time evolution through Langevin dynamics simulation using a friction coefficient, $\gamma$ = 0.05 at a temperature, T= 300 K and a salt concentration of 120 mM. The protein molecule was allowed to nonspecifically scan the DNA using both electrostatic and excluded volume interactions. We performed 100μs long production runs (1X $10^8$ MD steps) during which the dynamics of the DNA Binding Protein to access the target DNA site was observed by varying the attractive strength ($\epsilon/k_BT$) of the crowder molecules. On reaching the target DNA site, a soft, attractive Lennard-Jones potential ensures the specific contacts formation between the protein recognition helix and DNA target base pairs.

We performed an all-atom simulation of the Sap-1 protein in presence of PEG600 crowders. The simulation was performed using the GROMACS molecular dynamics package with the OPLS force field in the presence of PEG600 crowders at 300K. We used a base time step of 2 fs. The neighbour search was performed using the Verlet Algorithm and the Particle Mesh Ewald (PME) method treated the long range electrostatic interactions. We used the TIP3P water model to perform the simulations.

### 3. Criteria for data analysis:

The protein is said to perform 1D diffusion if it either slides or hops along the DNA surface. Sliding is confirmed in a snapshot if at least 70% of the recognition helix is within the DNA major groove and the centre of mass of the recognition helix is within 8Å to the nearest DNA base pair with an orientation angle < $25^{o}$.The protein performs 3D diffusion when the recognition region of the protein is at least 25Å away from the nearest DNA base pair. Hopping is considered in a snapshot when Sap-1 doesn't perform either sliding or 3D diffusion and is typically positioned 8-15 Å away from the closest DNA base pair. We consider the protein to be performing floating dynamics if it is 15Å -25Å away from the DNA surface and doesn't satisfy any of the sliding criteria.

#### 3.1 Calculation of depletion region around DNA:

The depletion region ($l_d$) around the DNA molecule is calculated by measuring the distance between the centre of each DNA base pair and the crowder molecule nearest to it throughout the simulation.

#### 3.2 Calculation of ruggedness chemical potential landscape:

To calculate the roughness of the chemical energy landscape, we followed the method adopted by Putzel *et al*(8). Here, we divided our simulation box into small cubic cells of side 50Å where the diffusing particle (Sap-1 protein) can move from each cell to another. While diffusing through the cells, the protein feels an effective potential, which is equal to protein's excess chemical potential. Therefore, by calculating the standard deviation of the average of each cell's potential gives a measure of ruggedness of the potential energy landscape.

The ruggedness of the chemical potential landscape is calculated as,

$$\sigma[ln(p^{cell})] = \sqrt{\frac{1}{N_{cells}}\sum_i \left(ln(p_i^{cell}) - \overline{ln(p_i^{cell})}\right)^2} \quad (18)$$

Where, $p_i^{cell}$ is the probability of the protein to be present in cell $i$. $N_{cells}$ is the number of cubic cells present inside the simulation box.

### 4. Tables

**Table1: Masses and Radius used for DNA components:**

| | Mass (Da) | Radius(Å) |
|---|---|---|
| Phosphate | 94.97 | 2.25 |
| Sugar | 83.11 | 3.20 |
| Adenine(A) | 134.1 | 2.70 |
| Thymine(T) | 125.1 | 3.55 |
| Guanine(G) | 150.1 | 2.45 |
| Cytosine(C) | 110.1 | 3.20 |

**Table 2: Masses and Radius used for amino acids:**

| Amino acid (single-letter code) | Mass (Da) | Radius of $C_\alpha$ (Å) |
|---|---|---|
| Isoleucine (I) | 131.1 | 2.0 |
| Lysine(K) | 146.1 | 2.0 |
| Phenylalanine(F) | 165.2 | 2.0 |
| Threonine(T) | 119.1 | 2.0 |
| Tryptophan(W) | 204.2 | 2.0 |
| Valine(V) | 117.1 | 2.0 |
| Arginine(R) | 174.2 | 2.0 |
| Histidine (H) | 155.1 | 2.0 |
| Alanine(A) | 89.0 | 2.0 |
| Asparagine(N) | 132.1 | 2.0 |
| Leucine (L) | 131.1 | 2.0 |
| Methionine(M) | 149.2 | 2.0 |
| Aspartic Acid (D) | 133.1 | 2.0 |
| Cytosine (C) | 121.1 | 2.0 |

| | | |
|---|---|---|
| Glutamic acid (E) | 147.1 | 2.0 |
| Glutamine (Q) | 146.1 | 2.0 |
| Glysine (G) | 75.0 | 2.0 |
| Proline (P) | 115.1 | 2.0 |
| Serine (S) | 105.0 | 2.0 |
| Tyrosine(Y) | 181.1 | 2.0 |

**Table 3: List of various sizes of PEG crowders as obtained from Sabirov et al**(9).

| **PEG crowders** | **Hydrodynamic radius(Å)** |
|---|---|
| PEG600 | 7.8 |
| PEG1000 | 9.4 |
| PEG1450 | 10.5 |
| PEG2000 | 12.2 |
| PEG3000 | 14.4 |
| PEG3400 | 16.3 |
| PEG4600 | 21 |
| PEG6000 | 25 |
| PEG20000 | 51 |

### 5. Figures:

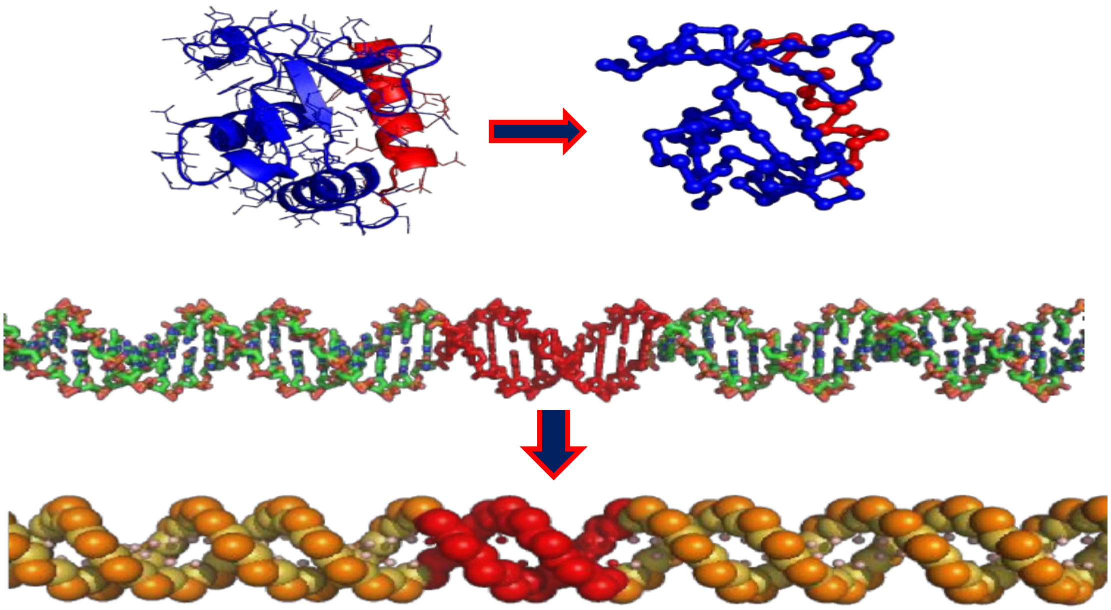

**Fig. S1.** Schematic representation of Sap-1 and B-DNA molecule in all atom (top left and middle) and coarse-grained (top right and bottom) models. The Sap-1 recognition helix is highlighted with red colour and corresponds to the residue number from 53 to 68 in the protein. Each nucleotide in the coarse grained model of DNA is presented by three beads at the centre of phosphate (orange colour beads), sugar (yellow colour beads) and base (pink colour beads). The 9 base pair DNA target site region is represented with red colour.

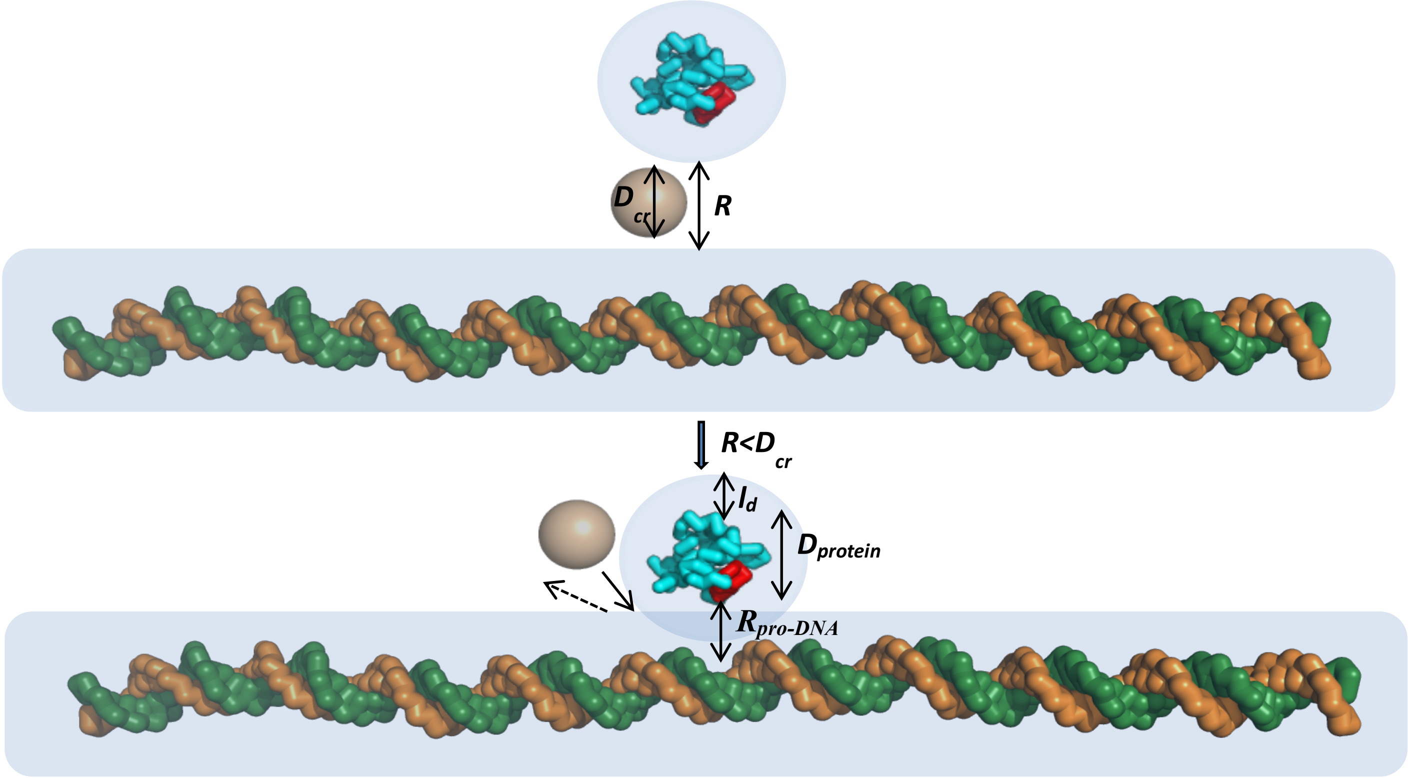


**Fig. S2**: Schematic representation of the origin of a composite depletion zone surrounding the protein and the DNA molecule in a crowded environment. When the distances between the surfaces of two molecules (R) get closer than the diameter of the crowder molecule ($D_{cr}$), the depletion zones surrounding the individual molecules merge to give rise to a composite depletion zone with decreased combined volume. This composite depletion zone comprises of the depletion region around the protein molecule ($l_d$), the diameter of the protein ($D_{protein}$), and the nearest distance between the protein and the DNA ($R_{pro\text{-}DNA}$) while the protein scans the DNA surface.

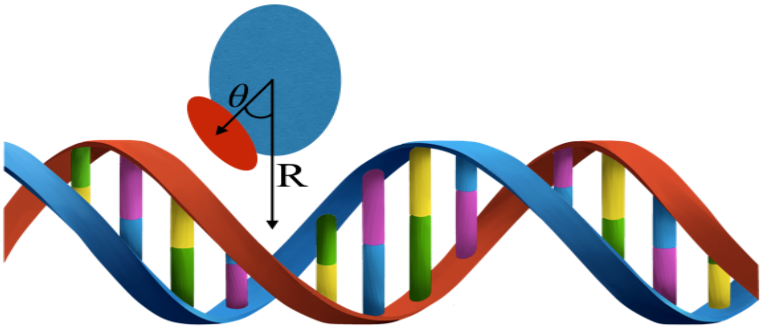


**Fig. S3 .** Schematic representation describing the various search modes (Sliding, Hopping and 3D Diffusion) adopted by the searching protein. The protein (blue coloured sphere with brown ellipse) is assumed to perform 3D diffusion if the centre of recognition helix (brown coloured ellipse) of the protein is at least 25 Å away (R > 25 Å) from the nearest DNA base pair. A snapshot is classified as sliding if ~70% of the protein recognition helix is in contact with the DNA major groove, R < 8 Å and the orientation angle (Θ) is <25°. A snapshot is classified as hopping if the protein doesn't perform either sliding or 3D diffusion.

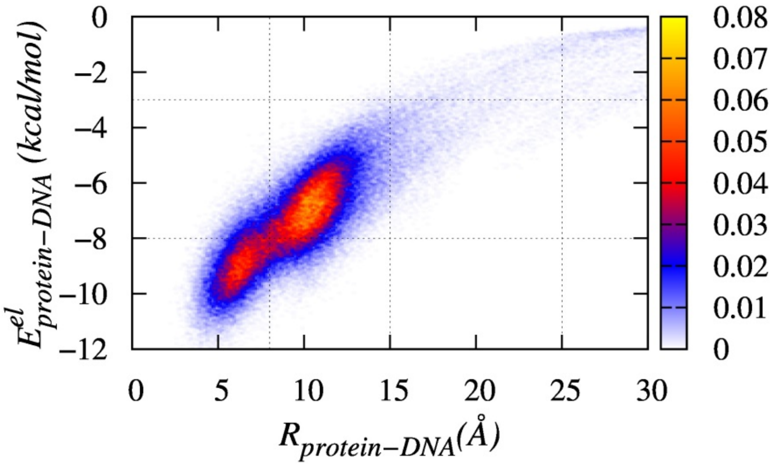


**Fig. S4**: Contour plot showing the relationship between the protein-DNA electrostatic interactions and the distance between the centre of recognition region of the protein and the nearest DNA base pair ($R_{protein\text{-}DNA}$). Here, the plot shows the probability of protein dissociation from the DNA surface ($R_{protein\text{-}DNA}$ >25 Å) in the absence of crowding agents.

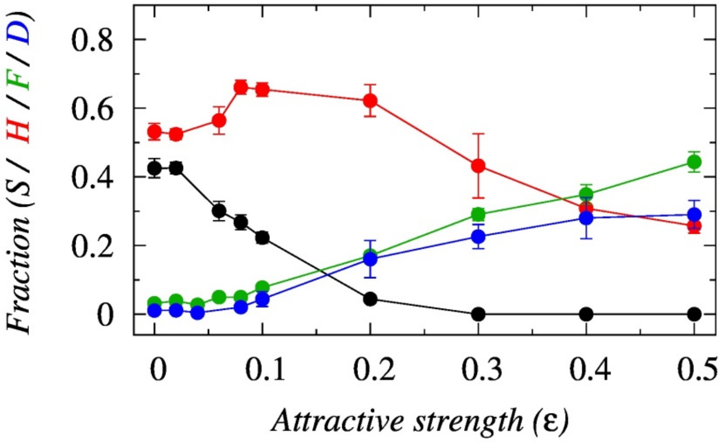


**Fig. S5:** Effects of crowder affinity (ϵ) on the target search mode of DBPs. (A) The variation in the affinities of different search modes (Sliding (S), Hopping (H), Floating (F) and 3D diffusion (D)) of the searching protein as a function of the ϵ.

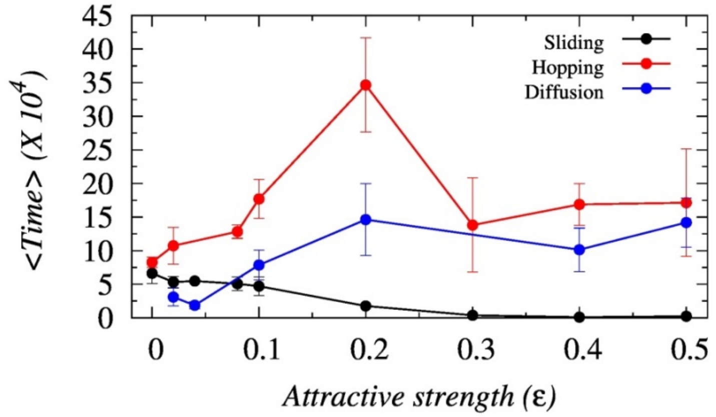


**Fig. S6** The variation in average duration spent by the searching protein performing sliding events (black line), hopping events (red line) and diffusion events (blue line) as a function of the interaction strength of crowder molecules (ϵ).

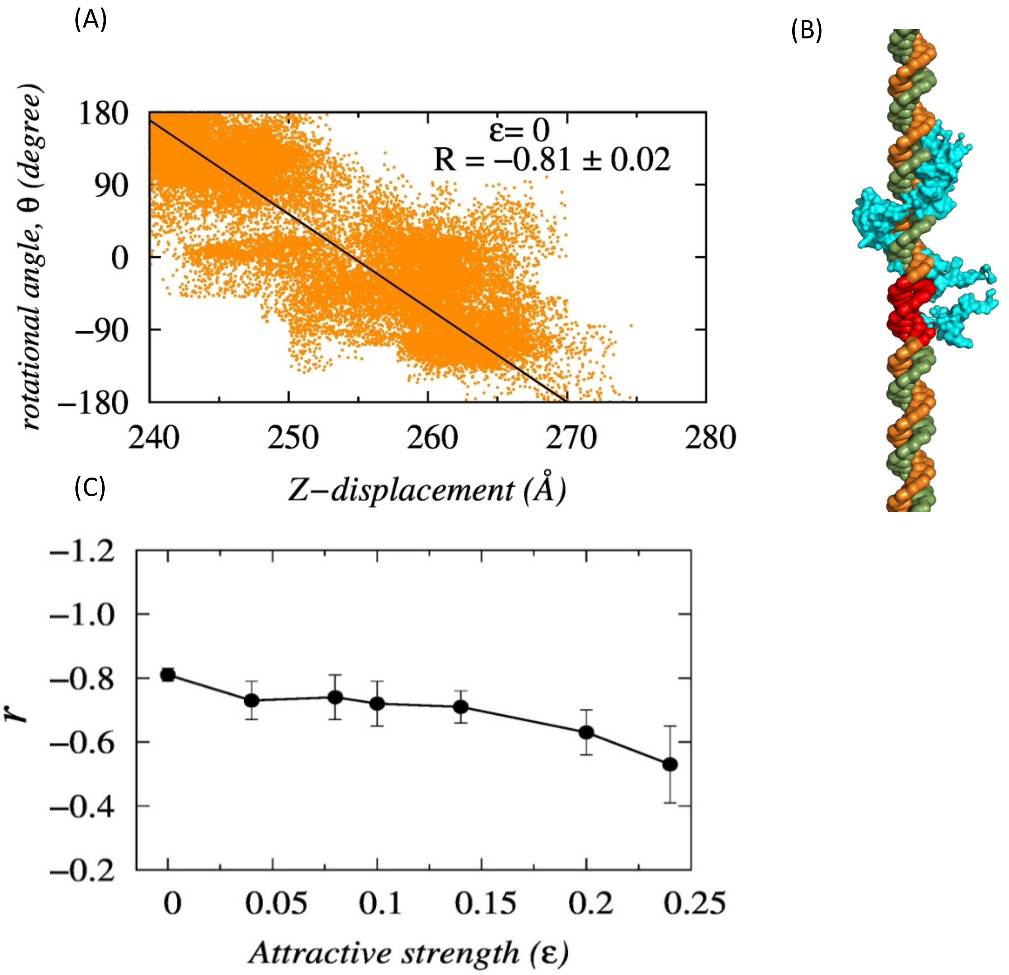


**Fig. S7 :** (A) The rotational coupled sliding motion performed by Sap-1 at $\epsilon$=0. Here, the correlation between the rotational motion of Sap-1 (θ) and its transversal displacement (Z-axis) is presented by the correlation coefficient, R. (B) Schematic representation of the rotational coupled sliding motion of Sap-1 while it scans the DNA by inserting its recognition helix into the DNA major groove. The target DNA site is shown in red. (C) Variation in the correlation coefficient (r) of the searching protein while performing rotationally coupled sliding along the DNA major groove with increasing $\epsilon$.

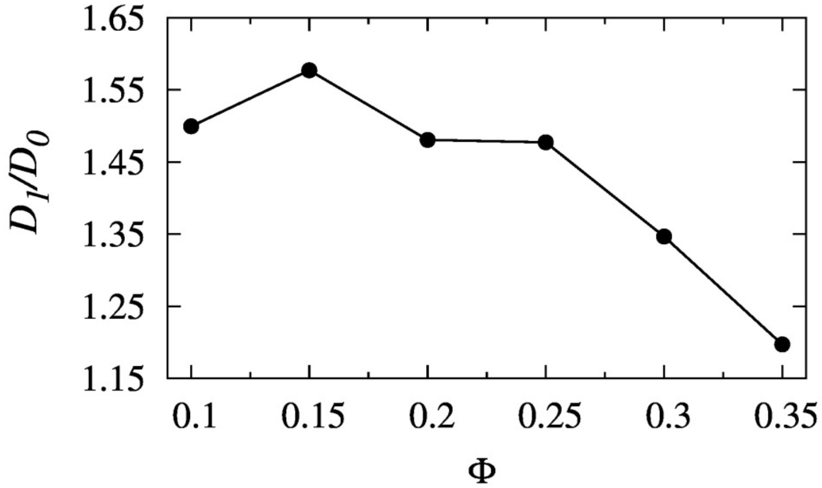


**Fig. S8:** Variation in the 1D diffusion coefficient of Sap-1 with increasing volume fraction of the crowder molecules (Φ). The diffusion coefficients are normalized by its value in the absence of crowder molecules.
